# Big trees drive forest structure patterns across a lowland Amazon regrowth gradient

**DOI:** 10.1101/2020.04.23.058289

**Authors:** Tassiana Maylla Fontoura Caron, Victor Juan Ulises Rodriguez Chuma, Alexander Arévalo Sandi, Darren Norris

**Author notes:** Corresponding author at: Coordenação de Ciências Ambientais, Universidade Federal do Amapá (UNIFAP), Rod. Juscelino Kubitschek Km 02, 68902-280 Macapá, AP, Brazil. Email address (D. Norris).

## Abstract

Degraded Amazonian forests can take decades to recover and the ecological results of natural regeneration are still uncertain. Here we use field data collected across 15 lowland Amazon smallholder properties to examine the relationships between forest structure, mammal diversity, regrowth type, regrowth age, topography and hydrography. Forest structure was quantified together with mammal diversity in 30 paired regrowth-control plots. Forest regrowth stage was classified into three groups: late second-regrowth, early second-regrowth and abandoned pasture. Basal area in regrowth plots remained less than half that recorded in control plots even after 20-25 years. Although basal area did increase in sequence from pasture, early to late-regrowth plots, there was a significant decline in basal area of late-regrowth control plots associated with a decline in the proportion of large trees. There was also contrasting support for different non-mutually exclusive hypotheses, with proportion of small trees (DBH <20cm) most strongly supported by topography (altitude and slope) whereas the proportion of large trees (DBH >60cm) supported by plot type and regrowth class. These findings support calls for increased efforts to actively conserve large trees to avoid retrogressive succession around edges of degraded Amazon forests.

## Introduction

Healthy tropical forests provide goods and services to human populations. Yet tropical forests show worrying rates of forest loss with an elevated loss / gain ratio and a statistically significant trend in annual forest loss of 2101 km^2^/year ^1^. One option to revert tropical forest loss is the restoration of degraded forests and deforested landscapes ^2,3^. Although the post-disturbance restoration of forest ecosystems often involves passive restoration strategies (i.e. natural regeneration), the ecological results of this type of restoration are still uncertain ^2–4^.

Continuing widespread forest losses across Amazonia compromises vital ecosystem services such as carbon storage, regulation of hydrological cycles and climate patterns ^5–7^. Riverside forests are particularly threatened and suffer losses due to the conversion of forest cover to pastures, compromising the maintenance of water flows ^8^. The recovery of degraded areas is necessary to recuperate the standing forest value and the Amazon offers an excellent recovery opportunity due to its natural potential for regeneration ^9,10^. Yet, the regrowth rate of degraded Amazon forests can be slow, as abandoned areas are typically on compacted poor quality soils ^11,12^ and due to the high structural and biological diversity of the original forests^13^.

Separating the complex interactions driving recruitment and recovery patterns of highly diverse Amazon forests is challenging ^2,3,14,15^, yet we know that different faunal groups can modulate and generate key impacts^16–19^. Indeed, the successional trajectory of natural regeneration in degraded forests can depend strongly on the concomitant recovery of faunal diversity and associated ecosystem services (e.g. seed dispersal) ^20–22^. For example, seed predation by both vertebrates and invertebrates ^23,24^ can limit germination and subsequent recruitment^22^. Long-term experiments have demonstrated the impact of vertebrates on recruitment, showing how this group contributes to the maintenance of tropical forest species and structural diversity ^25–28^.

Amazon mammals are important component of forest diversity ^25,29^ including carbon ^28^ and biomass cycles ^18^. Mammals can also play an important role in the successional trajectory and recovery of degraded areas as dispersers and predators of both seeds and seedlings ^23^. Mid- and large-bodied mammals (weight> 1 kg) can disperse a large numbers of seeds over long distances ^23,30^. For example, lowland tapirs can travel over 4 kilometers in a day ^31^ and disperse seeds of more than 70 tree species ^32^. The loss of mid- to large-bodied mammals may release some plant species from herbivory and increase their dominance, which subsequently decreases tropical forest biodiversity^33,34^.

Given the need to understand the patterns of forest structure in Amazonian forests, here we aim to identify how biotic and abiotic factors (Table 1) can explain patterns in forest structure across a successional gradient.

**Table 1.**
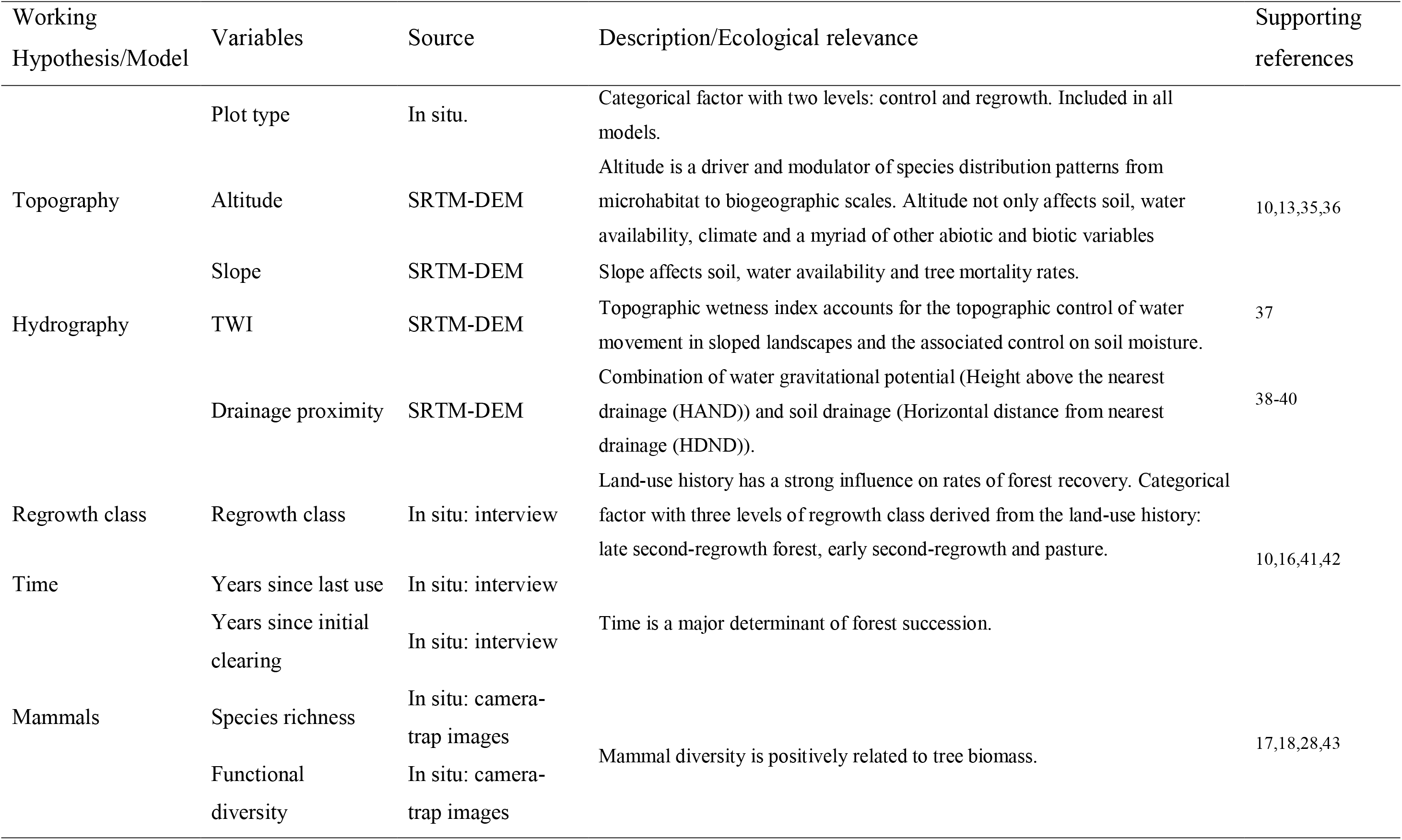
Explanatory variables.

## Results

### Variation in stand structure variables

There were clear differences in forest structure between control and regrowth plots (Figure 1). On average control plots had increased basal area and increased proportion of large trees (Figure 1). In contrast regrowth plots tended to have increased proportion of small trees (<20 cm DBH).

**Figure 1:**
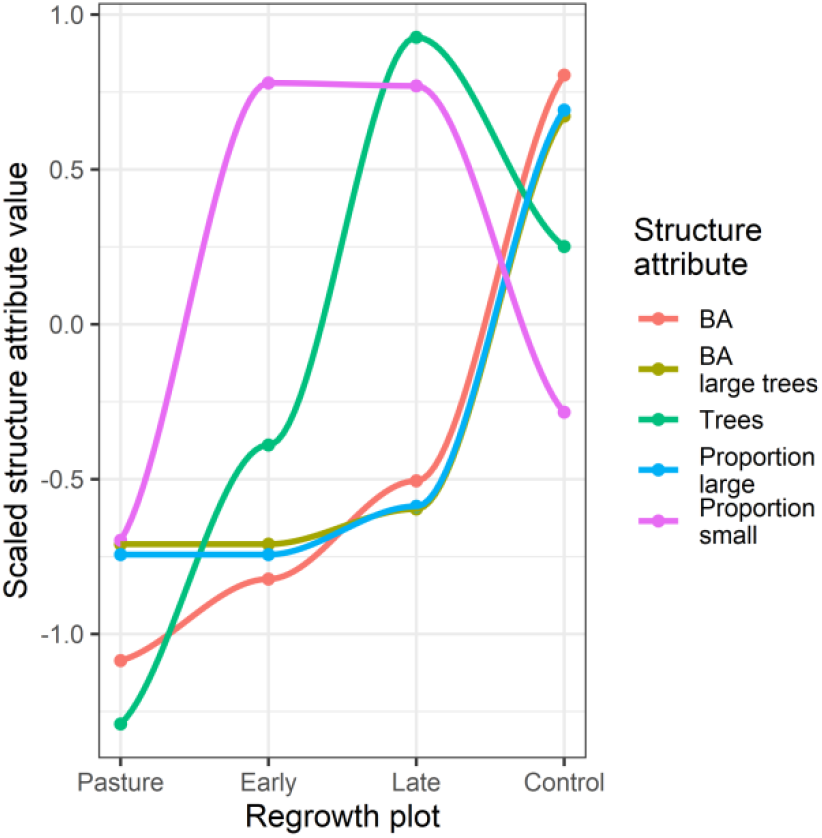
Forest structure changes across a lowland forest regrowth gradient. Showing mean values of five forest structure attributes recorded in 30 plots (15 control and 15 regrowth). Regrowth plot shows differences between control, late second-regrowth, early second-regrowth and pasture plots. Values are scaled (centered and scaled by the standard deviation) to enable simultaneous visual comparison of the different attributes. The lines are from LOESS smoothing as guides to aid visual interpretation.

The number and basal area of living trees tended to increase with altitude and this relationship was stronger in regrowth areas (Figure 2). The relationship with altitude was strongly affected by low lying (90 masl) pasture plots with no trees that generated significant leverage on the linear relationship (Figure 2).

**Figure 2:**
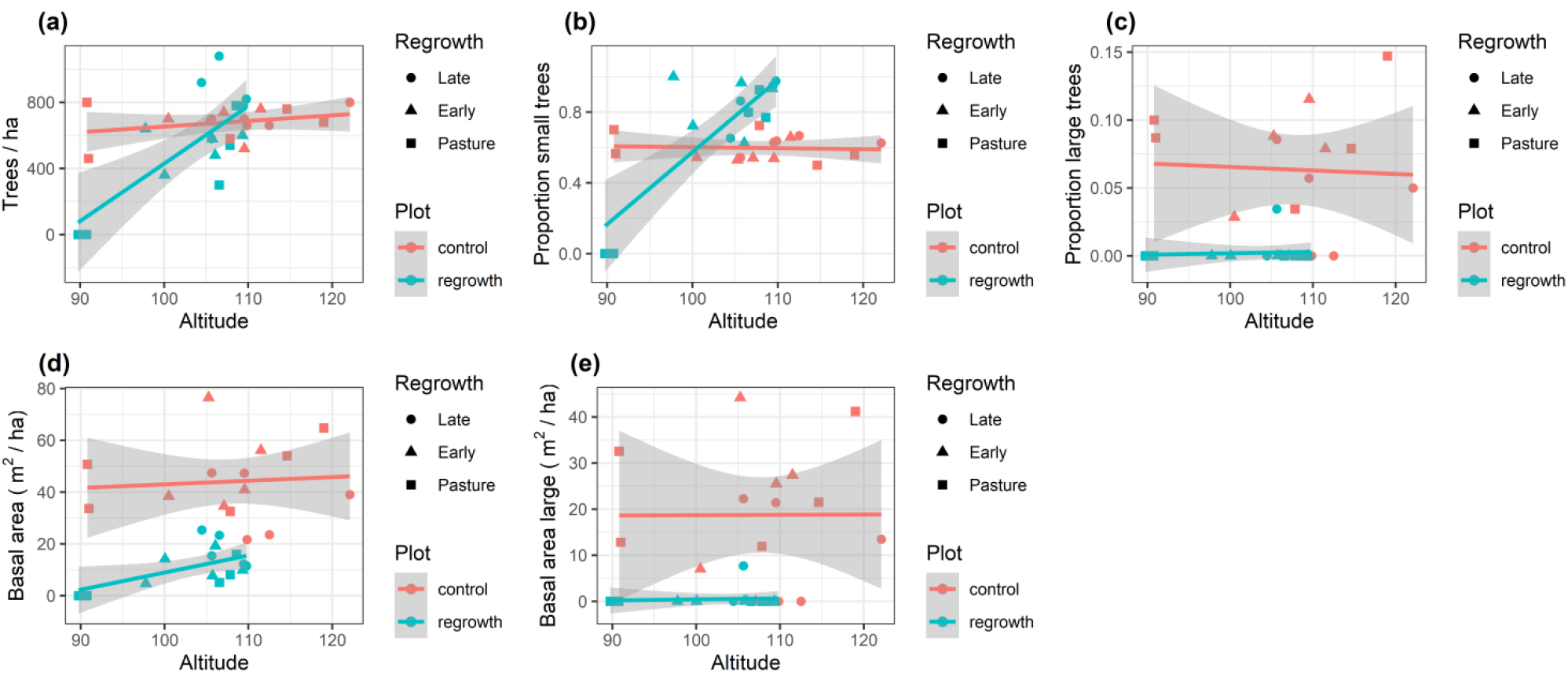
Forest structure along a lowland Amazon regrowth gradient. Showing trends in (a) number of trees (> 10 cm DBH) per ha, (b) proportion of small trees (10 – 20 cm DBH), (c) proportion of large (> 60 cm DBH) trees, (d) basal area and (e) basal area of large (>60 cm DBH) tree in 30 plots (15 control and 15 regrowth). Lines and shaded areas are mean values and 95% confidence intervals from linear models illustrating trends in basal area with increasing altitude (masl). Points with different shapes represent different regrowth classes.

Basal area ranged from 0 to 76.4 m^2^/ha across the 30 survey plots (Table 2), with control plots showing an average fourfold increase in basal area compared with regrowth plots (mean basal area 44.1 and 11.5 m^2^/ha, control and regrowth respectively, Figure 2). The patterns in plot basal area also differed between regrowth classes (Figure 3, Supplementary Table S1). There was a significant interaction between plot type (control/regrowth) and regrowth stage, with basal area increasing across pasture, early and late regrowth plots but control plots showing the opposite trend, with basal area decreasing significantly in late-regrowth control plots (Figure 3).

**Table 2.** Summary of survey locations. Characteristics of 15 sites used to study forest structure. Values are means with ranges in parentheses.

| Regrowth class | Sites/<br>Plots | Size<br>(ha) | Distance |  | Forest cover (%) |  | Tree BA<br>(m<sup>2</sup>/ha) |  | Large tree BA<br>(m<sup>2</sup>/ha) |  |
| --- | --- | --- | --- | --- | --- | --- | --- | --- | --- | --- |
|  |  |  | River<br>(m) | Town<br>(km) | 1 km | 5 km | Reg. | Cont. | Reg. | Cont. |
| Late | 5/10 | 5.8 | 288.15 | 40.4 | 91.5 | 97.4 | 17.6 | 35.8 | 1.5 | 11.4 |
|  |  | (2.0– | (110– | (35.0– | (87.4– | (96.4– | (11.5– | (21.7– | (0.0– | (0.0– |
|  |  | 12.0) | 554) | 45.0 | 95.4) | 98.5) | 25.3) | 47.4) | 7.7) | 22.3) |
| Early | 5/10 | 2.4 | 348.5 | 38.2 | 90.7 | 97.2 | 11.1 | 49.3 | 0.0 | 20.8 |
|  |  | (1.0– | (150– | (30.0– | (87.1– | (96.4– | (4.7– | (34.6– | (0.0– | (0.0– |
|  |  | 4.5) | 554) | 43.7) | 96.0) | 98.4) | 19.2) | 76.5) | 0.0) | 44.2) |
| Pasture | 5/10 | 8.7 | 266.8 | 32.8 | 88.8 | 96.9 | 5.8 | 47.1 | 0.0 | 24.0 |
|  |  | (6.8– | (170– | (26.8– | (85.7– | (95.5– | (0.0– | (32.6– | (0.0– | (12.0– |
|  |  | 9.9) | 461) | 40.8) | 91.1) | 98.5) | 15.9) | 64.8) | 0.0) | 41.2) |
| Totals | 15/30 | 5.6 | 301.1 | 37.1 | 90.4 | 97.2 | 11.5 | 44.1 | 0.5 | 18.8 |
|  |  | (1.0– | (110– | (26.8– | (85.7– | (95.5– | (0.0– | (21.7– | (0.0– | (0.0– |
|  |  | 12.0) | 554) | 45.0) | 96.0) | 98.5) | 25.3) | 76.4) | 7.7) | 44.2) |

**Figure 3:**
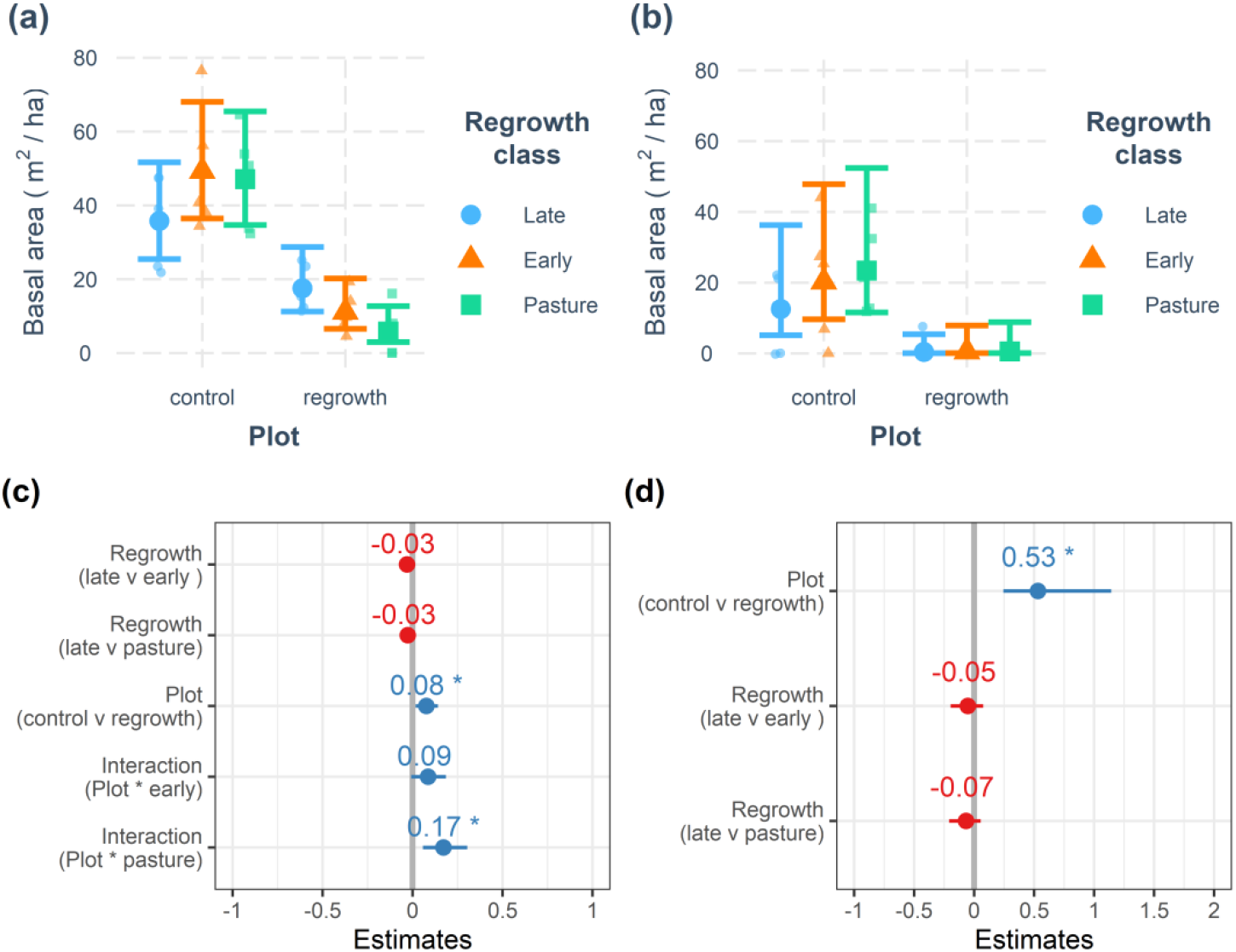
Basal area changes across a lowland forest regrowth gradient. The basal area of all (a and c) and large (b and d) living trees were recorded in 30 plots (15 control and 15 regrowth). Regrowth class shows differences between late second-regrowth, early second-regrowth and pasture plots contrasted with control forest plots. Top row shows Generalized Linear Model (GLM) predictions (mean and 95% confidence intervals) for basal area of (a) all and (b) large trees. Bottom row is the associated Forest-plot of the most parsimonious GLMs testing for interactions between regrowth class, plot type and years since last use in the basal area of (c) all and (d) large trees. Forest-plots show coefficient estimates and standard errors.

There was a highly significant positive linear relationship between overall basal area and large tree basal area (F_1,28_ = 127.5, R^2^ = 0.82, P < 0.0001). The basal area of large trees decreased significantly in regrowth compared with control plots (Figure 3). On average large trees accounted for 42% of the basal area in control plots compared with only 4% in regrowth plots (Table 2). Indeed a single large tree (>60 cm DBH) was recorded only once in a late-regrowth plot. This relationship was also reflected in the decline in basal area of late-regrowth control plots (Figure 3), which was associated with a decline on the proportion of large trees that accounted for a reduced 31% of the basal area in late-regrowth control plots (Table 2).

### Relationships between forest structure, mammal diversity and environmental variables

Mammal diversity varied considerably across the survey plots (Figure 4). There appeared to be a tendency for basal area to increase with mammal diversity in Late-regrowth plots, yet basal area was only weakly associated with mammal diversity within the different regrowth classes (Figure 4). Indeed, the diversity of mammals was found to be only weakly informative for explaining the basal area of trees across the 30 sample plots (Table 3).

**Table 3.** Summary of the Generalized Linear Models created to explain forest structure in 30 plots (15 control and 15 regrowth). Models ordered by decreasing AIC (Akaike Information Criterion) values.

| Forest structure | Model<sup>a</sup> | Dev. Exp | Loglik | BIC | AIC | Δ AIC | W<sub>i</sub> AIC<sup>b</sup> |
| --- | --- | --- | --- | --- | --- | --- | --- |
| Trees |  |  |  |  |  |  |  |
|  | Topography | 60.5 | -100.6 | 231.9 | 219.3 | 0.0 | 0.91 |
|  | Regrowth class | 49.8 | -105.1 | 234.1 | 224.3 | 5.0 | 0.07 |
|  | Mammals | 51.3 | -104.7 | 240.0 | 227.3 | 8.1 | 0.02 |
|  | Hydrography | 45.8 | -106.3 | 243.1 | 230.5 | 11.3 | 0.00 |
|  | Time | 42.9 | -107.0 | 244.7 | 232.1 | 12.8 | 0.00 |
|  | Plot | 6.5 | -114.4 | 239.1 | 234.9 | 15.6 | 0.00 |
| Proportion small trees |  |  |  |  |  |  |  |
|  | Topography | 75.2 | -64.3 | 155.7 | 144.5 | 0.0 | 0.96 |
|  | Hydrography | 70.4 | -67.5 | 162.2 | 151.0 | 6.5 | 0.04 |
|  | Mammals | 66.4 | -70.2 | 167.6 | 156.4 | 11.9 | 0.00 |
|  | Time | 65.1 | -71.0 | 169.3 | 158.1 | 13.6 | 0.00 |
|  | Plot | 53.7 | -78.7 | 164.3 | 161.5 | 17.0 | 0.00 |
|  | Regrowth class | 55.4 | -77.6 | 175.6 | 167.2 | 22.7 | 0.00 |
| Proportion large trees |  |  |  |  |  |  |  |
|  | Plot | 54.9 | -30.5 | 67.8 | 65.0 | 0.0 | 0.88 |
|  | Regrowth class | 60.6 | -28.9 | 78.2 | 69.8 | 4.8 | 0.08 |
|  | Mammals | 62.9 | -28.3 | 83.7 | 72.5 | 2.8 | 0.02 |
|  | Time | 62.8 | -28.3 | 83.8 | 72.5 | 2.8 | 0.02 |
|  | Topography | 54.7 | -30.6 | 88.5 | 77.3 | 7.5 | 0.00 |
|  | Hydrography | 52.8 | -31.1 | 89.5 | 78.3 | 8.5 | 0.00 |
| BA all trees |  |  |  |  |  |  |  |
|  | Topography | 80.7 | -100.9 | 232.3 | 220.4 | 0.0 | 0.89 |
|  | Regrowth class | 72.9 | -105.8 | 235.4 | 225.6 | 5.2 | 0.06 |
|  | Plot | 62.9 | -110.4 | 231.0 | 226.8 | 6.4 | 0.04 |
|  | Mammals | 71.7 | -106.4 | 243.4 | 230.8 | 10.4 | 0.01 |
|  | Time | 69.3 | -107.6 | 245.8 | 233.2 | 12.8 | 0.00 |
|  | Hydrography | 66.0 | -109.1 | 248.8 | 236.2 | 15.9 | 0.00 |
| BA large trees |  |  |  |  |  |  |  |
|  | Plot | 58.3 | -61.5 | 133.1 | 128.9 | 0.0 | 0.87 |
|  | Regrowth class | 67.1 | -59.9 | 143.5 | 133.7 | 4.8 | 0.08 |
|  | Hydrography | 70.9 | -59.0 | 148.7 | 136.1 | 7.1 | 0.02 |
|  | Mammals | 70.4 | -59.2 | 148.9 | 136.3 | 7.4 | 0.02 |
|  | Time | 65.7 | -60.1 | 150.9 | 138.3 | 9.4 | 0.01 |
|  | Topography | 60.8 | -61.0 | 152.7 | 140.1 | 11.2 | 0.00 |

<sup>a</sup> Models used to explain forest structure. All models contained plot type (control/regrowth) as categorical factor. Variables and associated estimates in the different models can be found as Supplementary Information Table S2 online.
<sup>b</sup> Akaike weights (W<sub>i</sub>) from largest to smallest.

**Figure 4.**
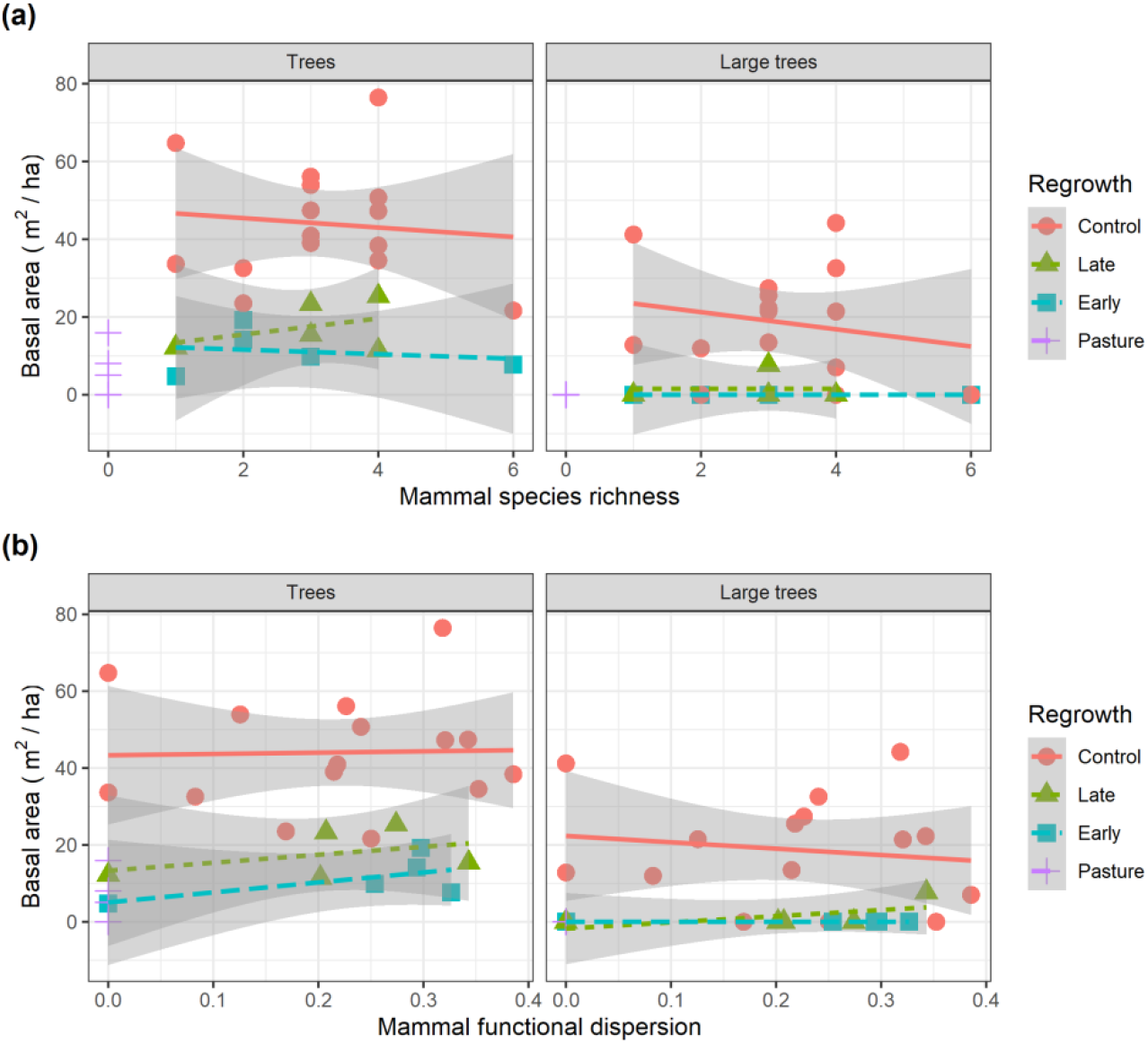
Mammal diversity and basal area across a lowland forest regrowth gradient. The basal area of (a) all and (b) large living trees were recorded together with the diversity (species richness and functional dispersion) of terrestrial mammals in 30 plots (15 control and 15 regrowth). Lines and shaded areas are mean values and 95% confidence intervals from linear models illustrating trends in basal area with increasing mammal diversity. Points with different shapes represent different regrowth plot types.

Comparison of the models representing the alternative hypothesizes showed that plot type (control v regrowth) and topography were the most important first ranked variables for the five forest structure attributes (Table 3). The most simple model including only plot type explained more than 50% of model deviance for all forest structure attributes except for the number of trees (DBH>10 cm). Plot type and regrowth class were both included in the 95% confidence set of models for the basal area of large trees (Table 3). In contrast Topography was the most important (first ranked) model for the number of trees, proportion of small trees and tree basal area (Table 3). Mammal diversity, Time and Hydrography models were not well supported and were not included in the 95% confidence set of models for any of the forest structure attributes (Table 3).

## Discussion

We integrate field and remotely sensed data to establish support for multiple non-mutually exclusive hypotheses explaining patterns in forest structure across a lowland Amazon regrowth gradient. We establish that different hypotheses are supported for different structure attributes. Here we discuss these findings in terms of prospects for the passive restoration of degraded Amazon forests.

The mean basal area value from our 15 control plots (44.1 m^2^/ha) was close to the mean from 42 Guyana Shield forest plots (43.4 m^2^/ha, range 10 – 65 m^2^/ha) in French Guiana ^35^. The results from Molto, et al. ^35^ were obtained from an extensive survey of 0.5 – 1 ha plots. Although our plot size was smaller compared to Molto, et al. ^35^, the similarity in mean values suggests that our plots do provide a representative sample of forest structure in the regrowth areas. The basal areas obtained from our regrowth plots followed a similar trajectory to those reported from abandoned pasture in Costa Rica ^10^, where the most recently abandoned pasture plots (<14 year) had mean basal area of 13.5 m^2^/ha, with basal area increasing to 26.1 m^2^/ha after 21 – 30 year ^10^.compared with 11.1 and 17.6 m^2^/ha respectively in our Early (1-5 year) and Late (20 – 25 year) regrowth plots. This also follows a similar pattern to values reported from 370 successional forest plots in the Brazilian Amazon, with basal area values typically < 10 m^2^/ha in early stages (< 5 year) and reaching 25 m^2^/ha after 15 years ^44^.

Although results from lowland forest sites in Costa Rica suggest rapid recovery of pasture areas ^10^ this could be related to the substantially lower basal area in the seven old growth reference plots (26.1 m^2^/ha, range 19.3 – 32.2 m^2^/ha) compared with those in our study area. Our results are similar to those reported from the central Amazon, where 25 y of regrowth restored half of the mature-forest biomass ^41^. A recent analysis of 45 Neotropical secondary forest study sites found that secondary forests in the lowland tropics reach 90 percent of old growth biomass in a median time of 66 yr ^13^. Our findings do suggest nuanced difference in successional trajectories. Basal area increased rapidly in early regrowth stages and this could be explained by the less intensive land use (i.e. lack of pasture) and the proximity to large areas of intact forest. In contrast basal area of late-regrowth areas was less than those reported from other areas ^10,44^. This could be related to soil productivity, as previous studies show that highly diverse Guyana Shield wet forests can take longer to establish ^13^. With basal area of control plots dominated by large trees it seems likely that many decades will be necessary for forest structure (total basal area, proportion of large trees) to return to pre-disturbance values.

The success of active and passive restoration can depend on ecological conditions ^45^. We found topography was the most informative model for explaining patterns in number of tress, tree basal area and proportion of small trees (Table 3, Figure 2). Differences in altitude and slope have been shown to affect floristic structure of tropical forests from local to regional scales ^16,35,42,46–48^. Indeed, even relatively small variations in topography can generate changes in local scale soil chemistry, hydrology and microclimate ^46,49^. The effects of topography do not operate in isolation from hydrology and the increased numbers of small trees and tree biomass with increasing altitude (Figure 3) agree with previous studies that show trees grow more slowly in more low lying (and often more waterlogged) terrain ^42^.

We found a weak association between mammal diversity and regrowth forest structure. Previous studies in a nearby protected area show that this group of mammals (mid–to large–bodied Artiodactyla, Perissodactyla and Rodentia) are more strongly associated with factors such as access to water ^50^ and altitude ^50,51^. A recent study also showed that mammal abundances were more strongly associated with phenology (fruit fall) than basal area along 10 km of forest in the western Guyana Shied ^52^. Additionally, regrowth class was found to be the primary driver of mammal species encountered independent of forest cover ^53^. For example the number of species detected in control and regrowth plots (all with forest cover >87%) varied between 1 and 6 (Figure 4). Mid- to large- bodied seed dispersers are a critical component of Amazon forests ^18,19,25^ and are also widespread and ubiquitous across myriad Amazonian forest types ^54–56^. The eight species are therefore not strictly dependent on the quality of forest habitat compared with other more specialist groups such as primates ^57^. The lack of a strong relationship between diversity of these eight mammal seed dispersers and forest structure attributes (i.e. overall basal area and proportion of small trees) is therefore to be expected.

Decades of research show that myriad edge effects can extend up to 150 m in fragmented Amazon forests ^58,59^. Considering the range of expect edge-effects it is highly probable that the natural regeneration and/or restoration of regrowth habitats in Amazon small-holdings (typically < 100 ha) will strongly depend on species ecological responses to habitat edges ^60^. Previous studies show that edge effects increase mortality of large trees, which in turn has major impacts on forest ecosystems ^61^. In highly fragmented areas edge-effects can drive tree communities through a process of “retrogressive succession”^62^ and toward an early successional state that may persist indefinitely. This early successional state can be characterized by functional and structural differences in that larger slower growing tree species with high wood density tend to decline whereas faster growing tree and liana species with lower wood density increase^62,63^. The decline in the number and basal area of large trees from our control plots along 20-25 year old edges suggest that retrogressive succession may establish even in relatively un-fragmented areas surrounded by extensive forest cover.

Our findings provide an early warning that even under a best case scenario there is potential for “retrogressive succession”. We found not only a lack of large trees in regrowth plots but also that large tree basal declined in older late-regrowth control plots. We suggest that this decline in large tress may be the primary driver of differences between regrowth and old growth forest and as such represent an unquantified component of resilience and time to recovery of Neotropical secondary forests. We also suggest that the continued presence of mid- and large bodied mammal seed dispersers in the study area are likely to be vital in order to avoid such “retrogressive succession”.

## Methods

### Ethics Statement

All methods were carried out in accordance with relevant guidelines and regulations. Fieldwork and data collection was conducted under research permit numbers SISBIO 40355–1, 47859-1 and 47859-2 to DN, issued by the Brazilian Ministério do Meio Ambiente (“MMA”). Data collection used non-invasive, remotely activated camera traps and did not involve direct contact or interaction with animals, thus no ethical approval was required. Interviews with local residents were approved by Brazilian Ministério do Meio Ambiente (SISBIO permits 45034-1, 45034-2, 45034-3) and the Ethics Committee in Research from the Federal University of Amapá (UNIFAP) (CAAE 42064815.5.0000.0003, Permit number 1.013.843). Interviews were conducted with residents that were both (1) willing to be interviewed (written informed consent was obtained from all interviewees) and (2) aware of the site history.

### Study area

Our study took place in 15 areas of regrowth on small holder properties^53,64^ in the center of the State of Amapá (Figure 5). The regional climate is classified by Koppen-Geiger as Am (Equatorial monsoon) ^65^, with annual rainfall greater than 2500 mm ^66^. The driest months are September to November (total monthly rainfall < 150 mm) and the wettest months from February to April (total monthly rainfall > 300 mm) ^66^. The State of Amapá has the lowest deforestation rate in Brazil and > 70% of the Amapá receives some form of legal protection. There is no large scale agricultural developments or monocultures along the waterways and properties retain typically small (< 1000 ha) areas of opened land, which are cleared for small scale family agriculture, which focuses on acai, small scale production of fruits and vegetables for sustenance and limited commercial sale of regional produce (e.g. manioc flour) in local markets. There are some 54 properties upstream of the nearest town (Porto Grande ^67^). There has never been any expansive clearcutting in the region and there are no monocultures (e.g. soy) or cattle production. All sites were at least 26 km from the nearest town by river, and all sites are surrounded by matrix of continuous closed canopy forest cover (Table 2). Pesticides and/or herbicides had never been used at any of the sites.

**Fig. 5.**
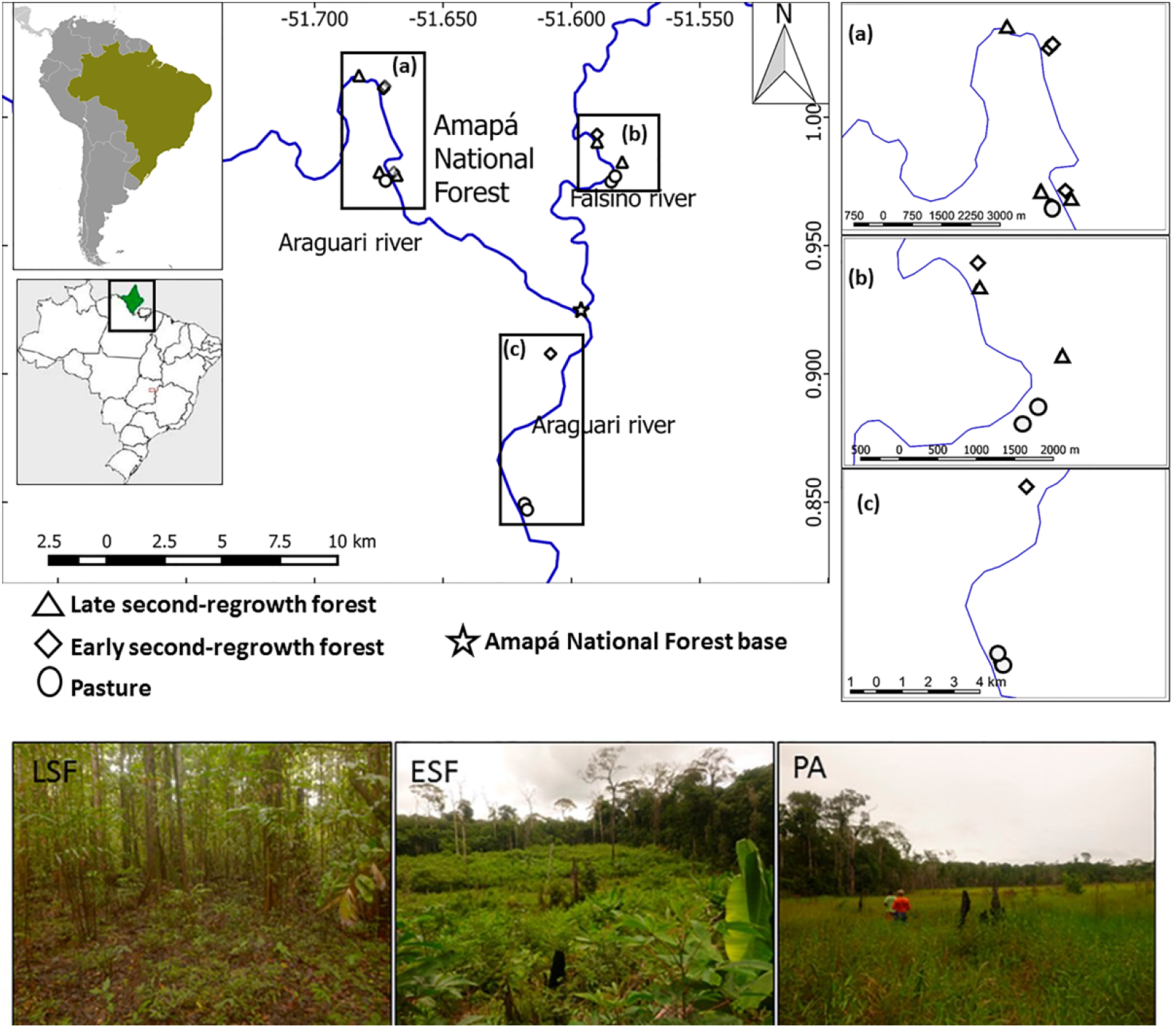
Map of the study area in the eastern Amazon. Showing the location of 15 study sites, grouped into three regrowth stages in the small holder properties close to rivers (solid blue lines): late second-regrowth forest (LSF, triangles), early second-regrowth forest (ESF, squares) and pasture (PA, circles).

A described previously^53,64^ the 15 small-holder properties were selected based on differences in land-use histories and forest succession/regrowth stage. All sites were close (110 – 554m, Table 2) to 100 – 200 m wide rivers that are navigable by motorized boats, but due to riverbank formation the sites are never flooded. These 15 sites were grouped into three regrowth classes based on the land-use history: late second-regrowth forest (N = 5, most recent human disturbance between 20 and 25 years), early second-regrowth (N = 5, most recent human disturbance between 1 and 5 years), and pasture (N = 5, recently cleared and abandoned pasture areas dominated by grasses/herbs but that had never been used to raise livestock, with the most recent disturbance between 1 and 17 years). Each of the 15 regrowth sites was paired with a nearby (60 to 150 m) control site i.e. 20 – 30 m tall *terra-firme* forest site without a history of mechanized timber extraction. To reduce the possible confounding influence of edge effects that are known to strongly influence the distribution of trees in Neotropical forests, all regrowth and control sites were established at a standardized distance (approximately 30 m) from the nearest control-regrowth habitat edge.

### Forest structure

Data were collected from May to August 2016. Forest structure data (i.e., number of trees and basal area) were obtained from plots measuring 50 x 10 m (500 m^2^), at each of the 30 points, totaling 1.5 hectare. This plot size was selected as it has been widely used to examine structural changes in tropical forests ^19,41,42,68^ and several of the regrowth areas were too small (Table 2) to enable the establishment of larger spatially independent plots. We obtained five measures (responses) to characterize the forest structure in each plot. These were selected based on previous studies that show their appropriateness to distinguish attributes of regrowth/successional stages related to biodiversity of Amazon forests ^13,44,69,70^. The number of all trees ≥ 10 cm DBH (diameter at breast height at a standard 1.3 m above ground, or above tallest root buttress) was used to quantify the number of trees per area in each plot (m^2^). This count included all trees which had at least half of their basal trunk inside the plot. The proportion of small (10 – 20 cm DBH) trees was calculated to represent the expected increase of younger trees in regrowth areas. The proportion of large (>60 cm DBH) trees was calculated as this is known as an important characteristic of mature/late succession areas ^44,70^. We also calculated the basal area of all and large trees as this is known to be strongly correlated with tree biomass ^71^. For example basal area and biomass were > 99% correlated in 23 plots from lowland Costa Rica ^10^.

### Explanatory variables

We investigated predictions from multiple non-mutually exclusive hypotheses to explain patterns in basal area (Table 1). A total of 10 variables were used to form models to represent 5 working hypotheses (topography, hydrography, regrowth class, time and mammal diversity) that based on the findings from previous studies were likely to explain the observed patterns ^10,13,28,40,41,72^. We chose to work with mainstream, widely available environmental variables. Four of these (the topographic and hydrographic model variables) were computed from remotely sensed digital terrain model (SRTM-DTM): altitude (masl), slope, TWI (Topographic wetness index), DND (Distance to Network Drainage) calculated from the interaction between HAND (Height above network drainage) and HDND (Horizontal distance to network drainage). The time model included years since the regrowth site was opened and years since last use, both of which were obtained from interviews with local landowners.

Mammal functional diversity was obtained from a camera-trap survey conducted at the same time (May to September 2016) and in the same plots as forest structure was sampled ^53^. Camera traps equipped with infrared triggers (Bushnell Trophy Cam, 8MP, Overland Park, KS, USA) were installed in each of the 30 plots following standardized protocols ^50,51,73^. This camera trap survey [full details provide in ^53^] including a sampling effort of 827 camera-trap days (450 and 377 camera-trap days, control and regrowth sites respectively) was used to estimate functional diversity of eight terrestrial mammal seed dispersers (*Cuniculus paca*, *Dasyprocta leporina*, *Myoprocta acouchy*, *Mazama americana*, *M. nemorivaga*, *Pecari tajacu*, *Tayassu pecari* and *Tapirus terrestris*).

### Data analysis

Tree Basal Area in each plot was obtained as the sum of the basal area value for each individual tree derived from the DBH of each tree following the formula BA (basal area in m^2^)= 0.00007854 X DBH^2^ (constant obtained by solving the following equation to obtain BA in m^2^ from the DBH measured in cm ^69^):

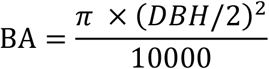

We calculated basal area of all and large (>60cm DBH) living trees ^69,70,72^. We also calculated the proportion of small stems (10 – 20 cm DBH trees) as this has been shown to be an important measure of stand structure in forest regrowth areas ^35,69^.

To represent diversity of terrestrial mammal seed dispersers we calculated a richness and functional diversity (FD) value for each of the 30 plots ^53^. Richness was calculated as the observed number of species (hereafter “species richness”) at each plot. Although there are many diversity metrics, we chose species richness as it is widely used and clearly interpretable ^74,75^ and with relatively few (eight) species and 30 plots there were strong correlations between species richness values and alternative diversity metrics such as Shannon and Simpson diversity (Spearman rho > 0.89). We used Functional Dispersion (FDis) ^76^ as an index of functional diversity as it is not strongly influenced by outliers, accounts for relative abundances, is unaffected by species richness and can be calculated from any distance/dissimilarity measure ^76,77^. Functional Dispersion was estimated with the dbFD function ^77^ using default settings.

To examine patterns in forest structure attributes we used Generalized Linear Models. We used an information theoretic model averaging framework ^78^ to examine the support for five models representing the five non-mutually exclusive hypotheses – topography, hydrography, regrowth class, time and mammal diversity (see Table 1 for variable description and ecological relevance). We evaluated models based on their information content, as measured by AIC – Akaike Information Criterion. The relative importance of the models was measured by the models Akaike weights (Burnham & Anderson 2002 pp. 75-77, 167-172), which is a scaled measure of the likelihood ratio that ranges between 0 (least important) and 1 (most important). None of the unexplained variation (model residuals) was related to the geographic distance among plots so we did not need to control for spatial dependence. All analysis were conducted using the R language and environment for statistical computing ^79^, with base functions and functions available in the following packages: vegan^80^, ggplot2 ^81^, MuMIn ^82^, and tweedie ^83^.

## Data Availability Statement

The raw forest structure and environmental data used in the analysis of this study have been deposited in the OSF - Center for Open Science at DOI: 10.17605/OSF.IO/MC27U.

## Acknowledgements

The Instituto Chico Mendes de Conservação da Biodiversidade (ICMBio) and the Amapá National Forest staff (Érico Emed Kauano and Sueli Gomes Pontes dos Santos) and the Federal University of Amapá (UNIFAP) provided logistical support. We thank the Brazilian Ministério do Meio Ambiente (“MMA”) for authorizing data collection (SISBIO permits 40355–1, 47859-1 and 47859-2). We also thank the local landowners who gave permission for data collection at their properties. We are deeply indebted to Cremilson and Cledinaldo Alves Marques and family for their dedication, commitment and assistance during the fieldwork.

## Author Contributions

D.N. conceived of the project; V.J.U.CR and A.A.S collected data. V.J.U.CR, A.A.S and D.N. performed data analysis and interpretation. A.A.S prepared figure 1. D.N. prepared figures 2–5. T.M.F.C and D.N. wrote the main manuscript text. All authors reviewed and revised the manuscript.

## Supporting Information

**Supplementary Table S1.** Generalized Linear Model values.

| *Predictors* | **All trees** |  |  |  |  |  | **Large trees** |  |  |  |  |  |
| --- | --- | --- | --- | --- | --- | --- | --- | --- | --- | --- | --- | --- |
|  | **Full** |  |  | **Best** |  |  | **Full** |  |  | **Best** |  |  |
|  | *Est.* | *CI* | *p* | *Est.* | *CI* | *p* | *Est.* | *CI* | *p* | *Est.* | *CI* | *p* |
| (Intercept) | 0.46 | 0.40–0.53 | **<0.001** | 0.46 | 0.43–0.50 | **<0.001** | 0.66 | 0.50–0.81 | **<0.001** | 0.64 | 0.55–0.76 | **<0.001** |
| Last use | 0.00 | -0.00–0.00 | 0.933 |  |  |  | -0.00 | -0.01–0.01 | 0.896 |  |  |  |
| Regrowth (late v early) | -0.01 | -0.10–0.07 | 0.826 | -0.03 | -0.08–0.01 | 0.176 | -0.04 | -0.24–0.17 | 0.710 | -0.05 | -0.20–0.08 | 0.441 |
| Regrowth (late v pasture) | -0.00 | -0.09–0.08 | 0.966 | -0.03 | -0.07–0.02 | 0.245 | -0.08 | -0.27–0.10 | 0.355 | -0.07 | -0.21–0.06 | 0.311 |
| Plot (control v regrowth) | 0.14 | 0.02–0.28 | **0.050** | 0.08 | 0.02–0.14 | **0.022** | 0.02 | -0.40–0.44 | 0.917 | 0.53 | 0.24–1.14 | **0.016** |
| Interaction (Last use * early) | -0.01 | -0.02–0.01 | 0.467 |  |  |  | -0.01 | -0.05–0.03 | 0.525 |  |  |  |
| Interaction (Last use * pasture) | -0.00 | -0.01–0.00 | 0.386 |  |  |  | -0.00 | -0.01–0.01 | 0.986 |  |  |  |
| Interaction (Last use * Plot) | -0.00 | -0.01–0.00 | 0.280 |  |  |  | 0.03 | -0.04–0.10 | 0.339 |  |  |  |
| Interaction (Plot * early) | 0.03 | -0.11–0.17 | 0.624 | 0.09 | -0.01–0.19 | 0.090 | 46.73 |  | 0.999 |  |  |  |
| Interaction (Plot * pasture) | 0.15 | 0.02–0.29 | **0.040** | 0.17 | 0.06–0.30 | **0.011** | 42.03 |  | 0.999 |  |  |  |
| Observations | 30 |  |  | 30 |  |  | 30 |  |  | 30 |  |  |
| R<sup>2</sup>Nagelkerke | 0.83 |  |  | 0.80 |  |  | 0.81 |  |  | 0.74 |  |  |
| AIC | 229.40 |  |  | 225.62 |  |  | 140.45 |  |  | 132.02 |  |  |

**Supplementary Table S2.**
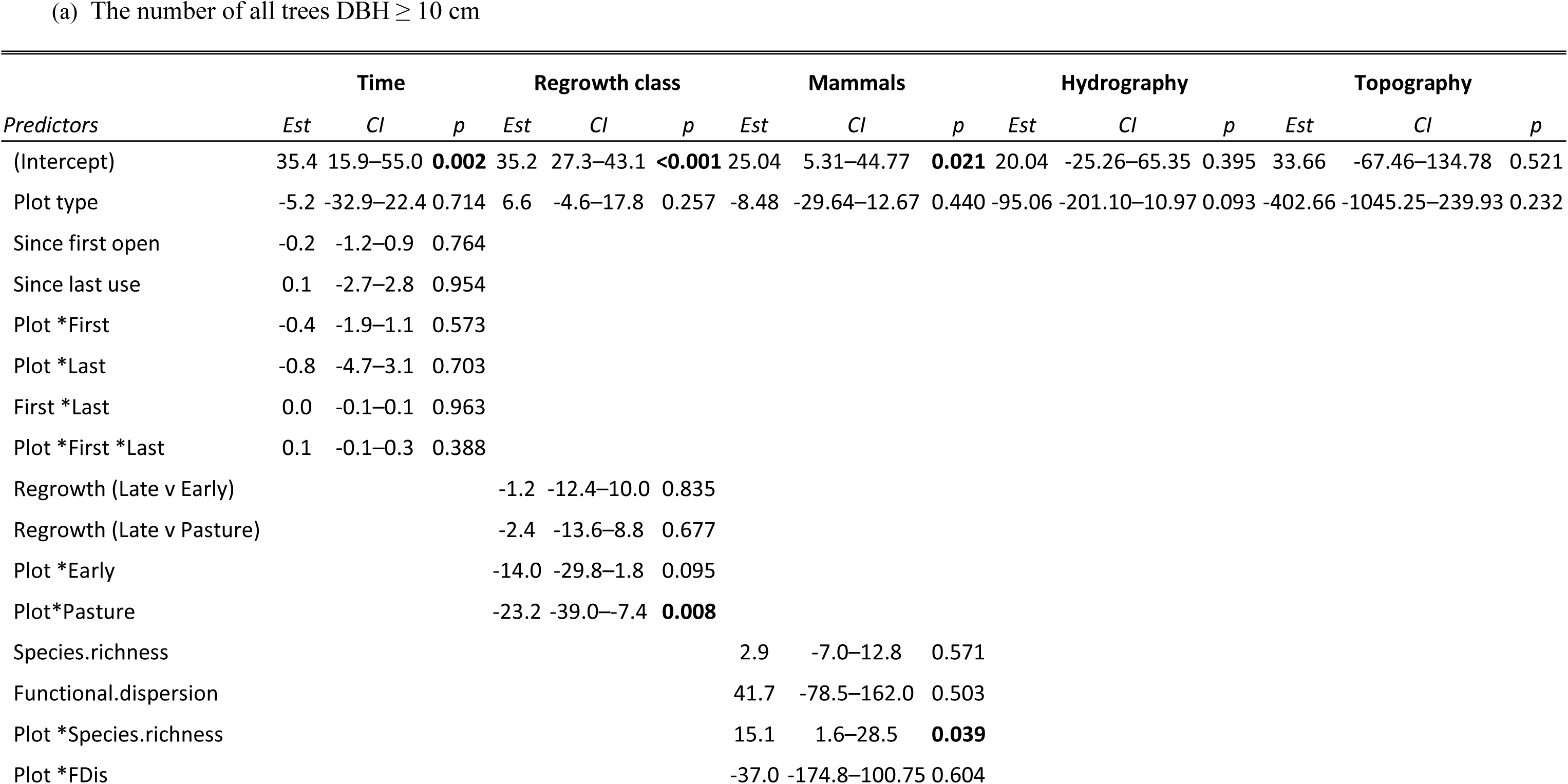
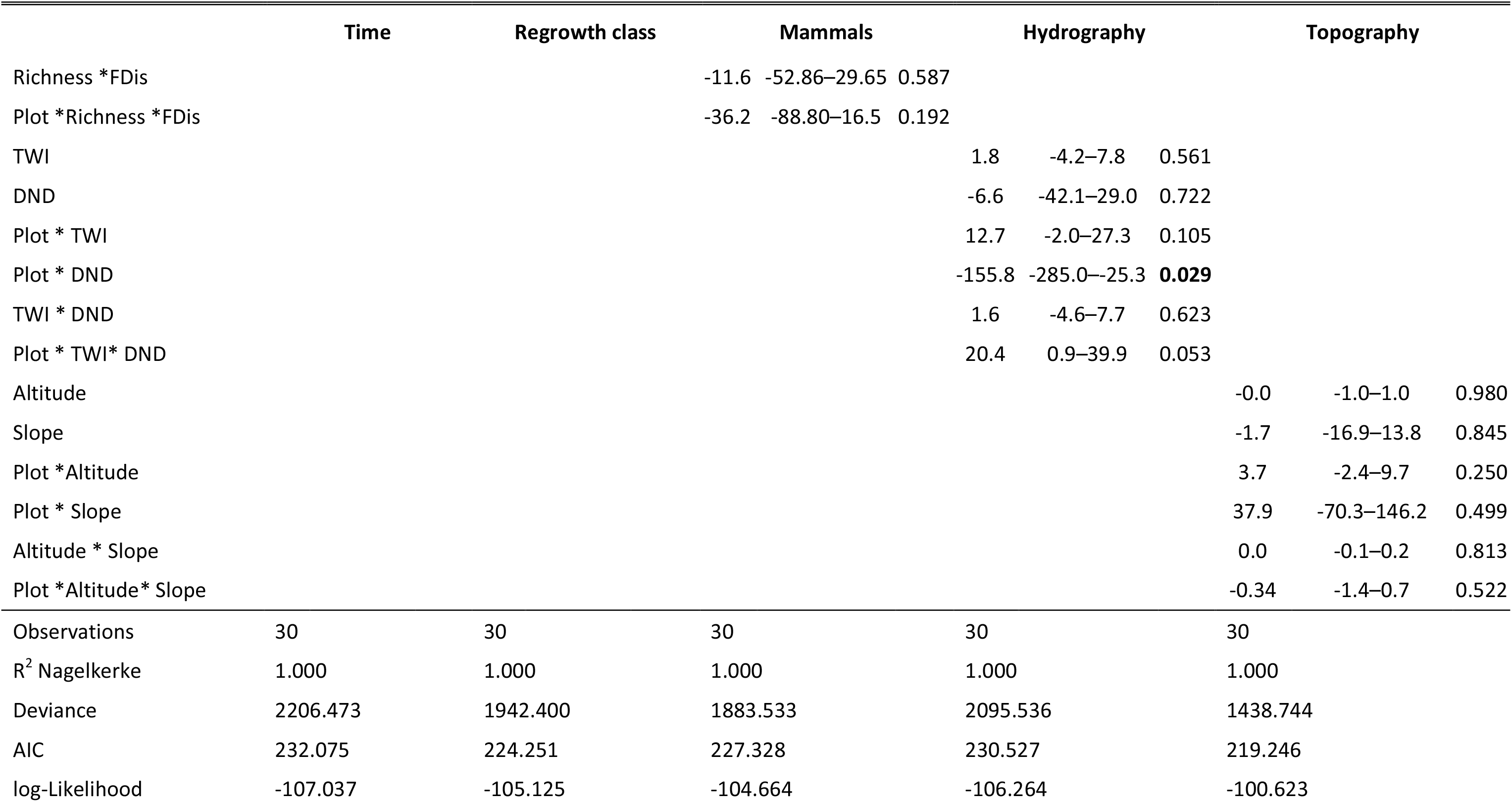
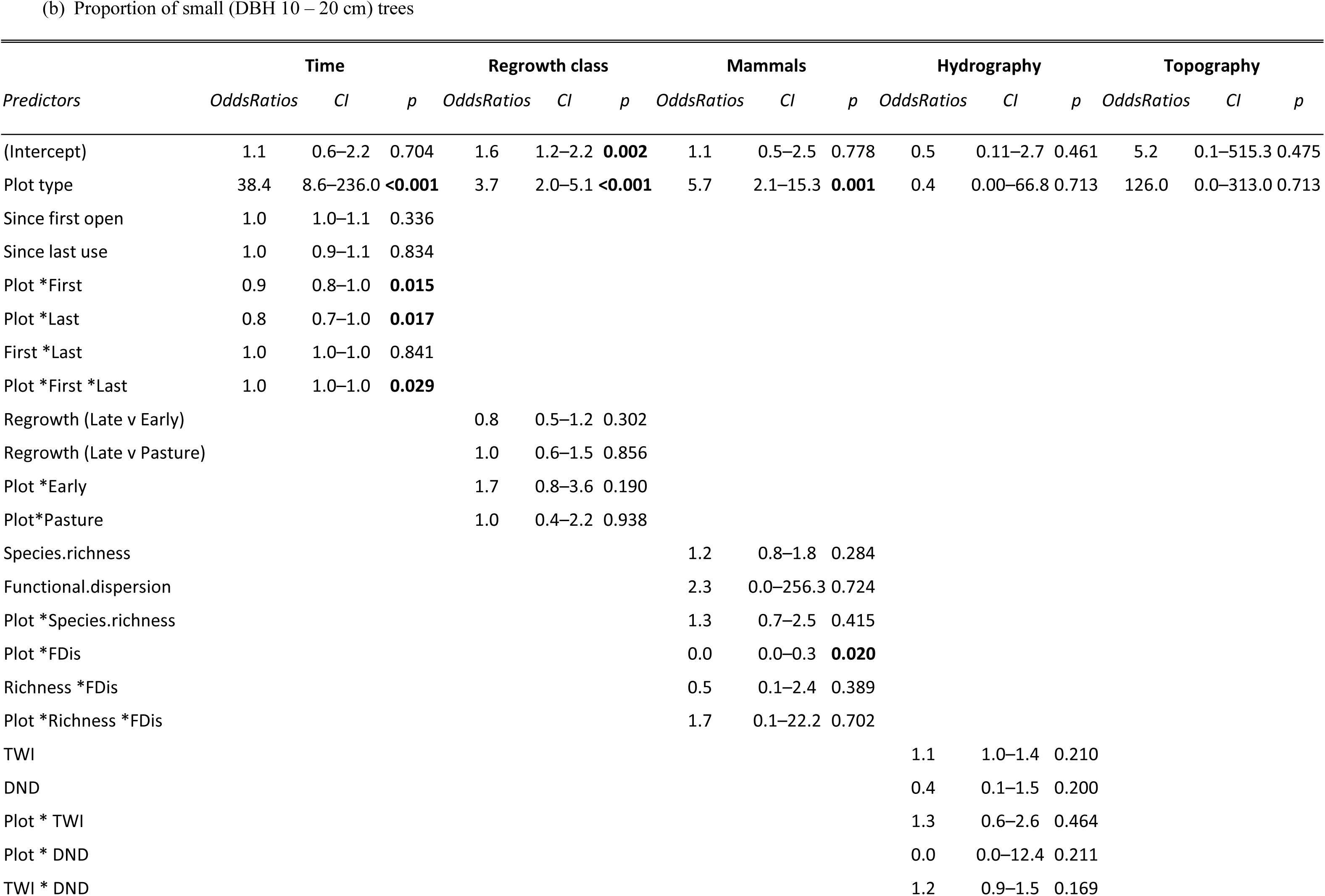
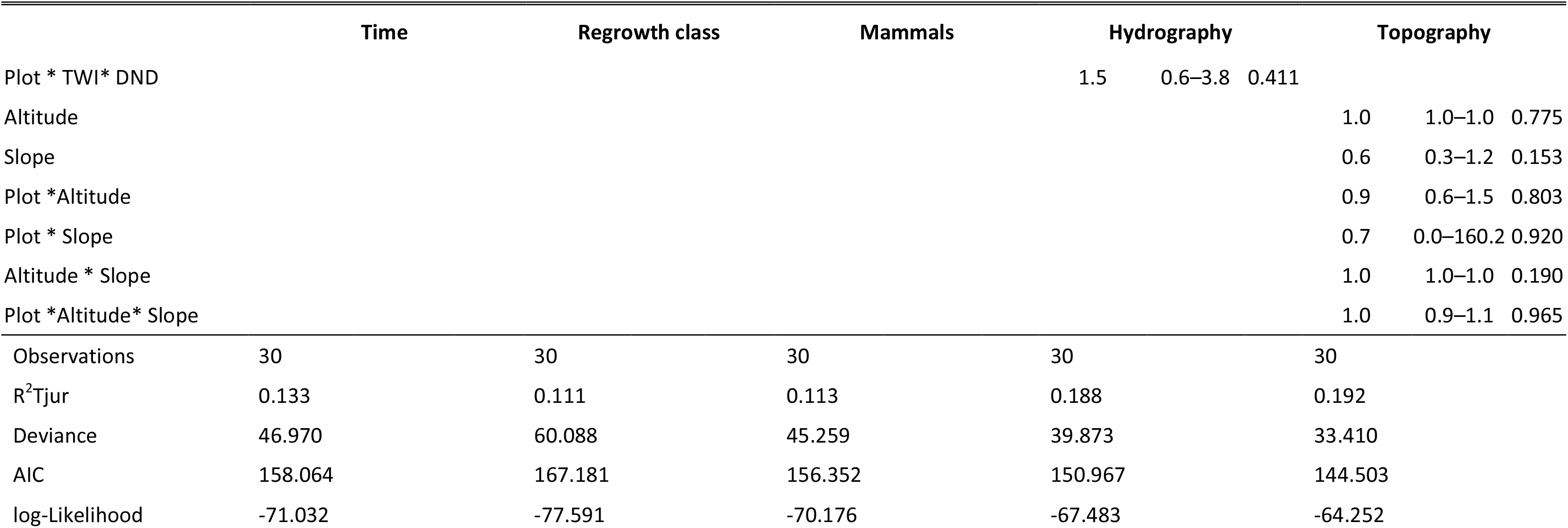
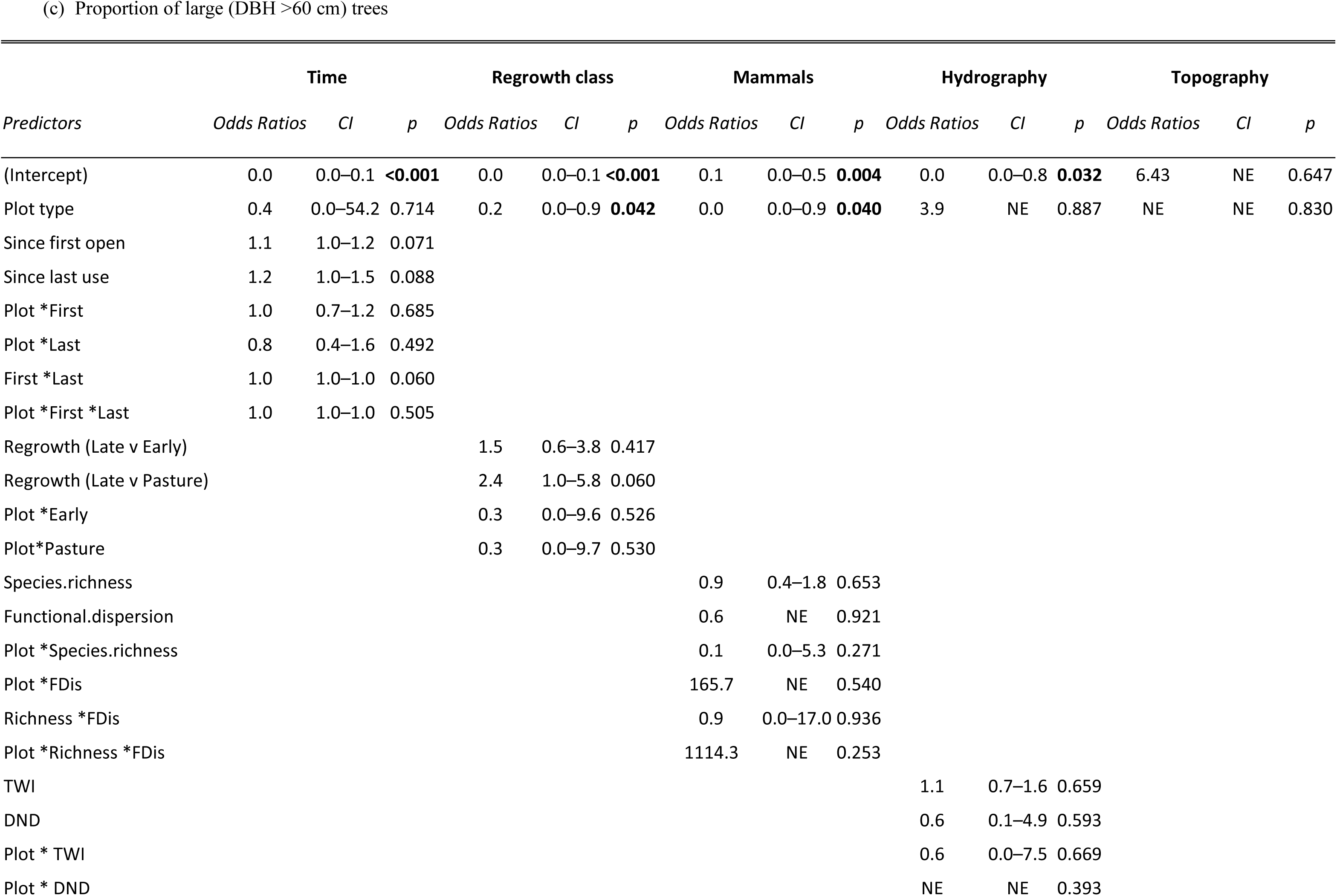
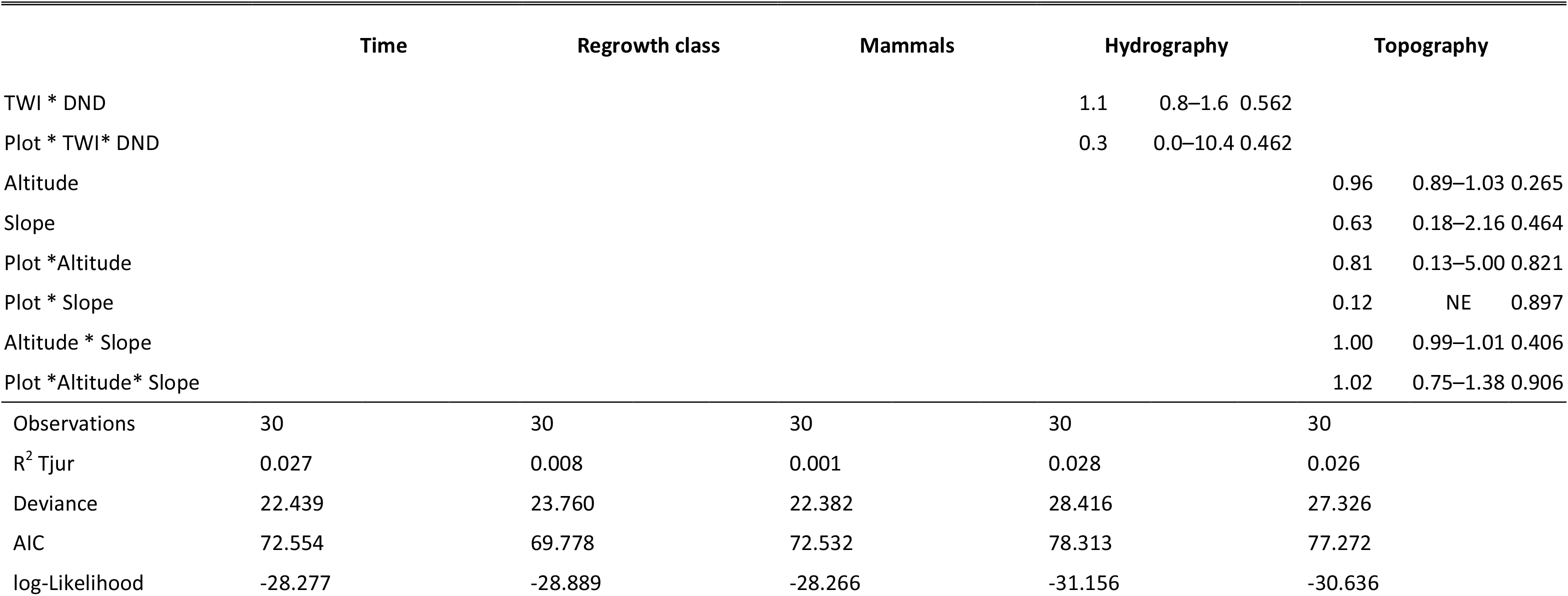
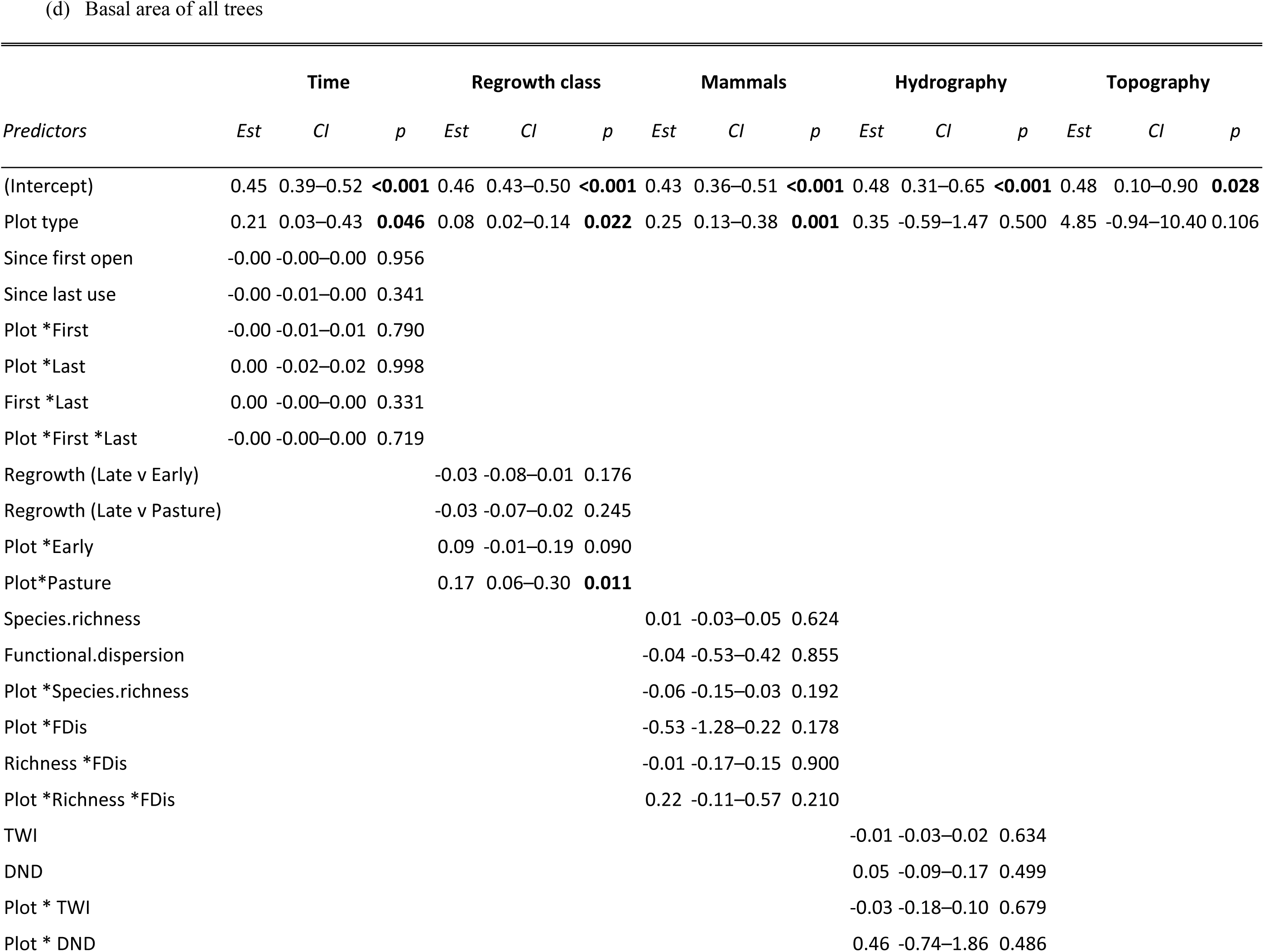
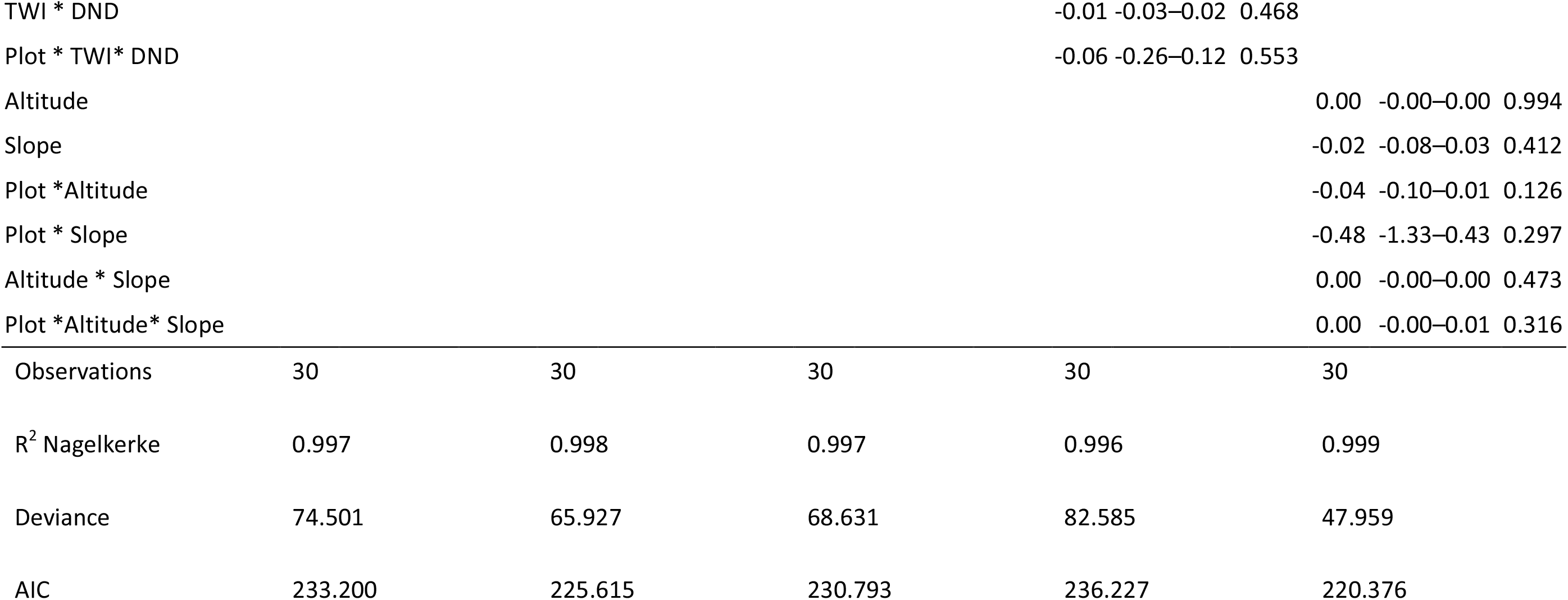
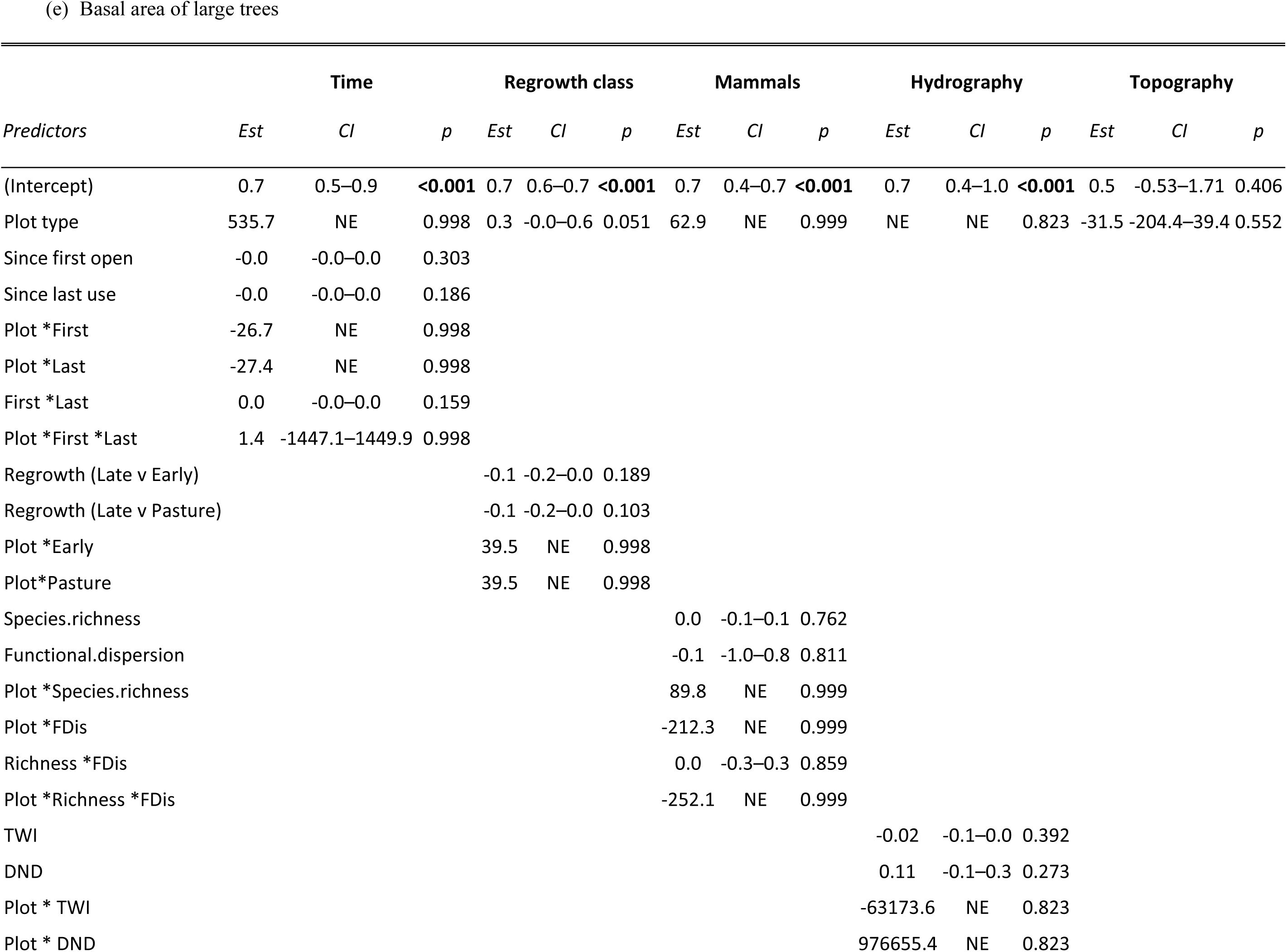
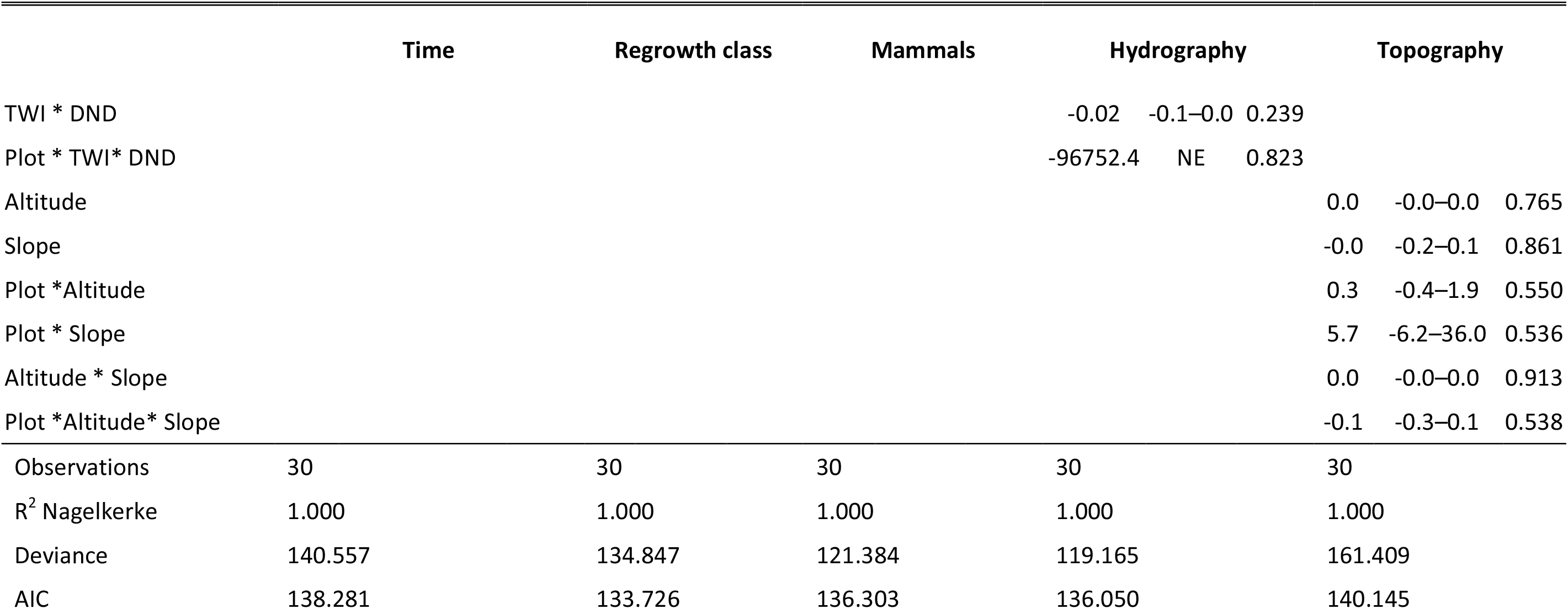
Generalized Linear Models created to explain forest structure in 30 plots (15 control and 15 regrowth). Model summaries for responses of five structural attributes: (a) The number of all trees ≥ 10 cm DBH; (b) Proportion of small (DBH 10 – 20 cm) trees; (c) Proportion of large (DBH >60 cm) trees; (d) Basal area of all trees; (e) Basal area of large trees. Showing slope estimates (“Est”) for 10 variables in 5 models (time, regrowth class, mammal diversity, hydrography and topography). “NE” denotes cases when values could not be reliably estimated. The time model included years since the regrowth site was opened and years since last use. The regrowth model included sites grouped into three regrowth classes (pasture, early-regrowth and late-regrowth). Topography included altitude (masl) and slope. Hydrography was modelled with TWI (Topographic wetness index), DND (Distance to Network Drainage) calculated from the interaction between HAND (Height above network drainage) and HDND (Horizontal distance to network drainage). Mammal diversity was obtained from camera-traps and quantified as species richness and Functional Dispersion (FDis).

